# Effects of progesterone on the reproductive physiology in zebrafish

**DOI:** 10.1101/147280

**Authors:** Chunyun Zhong, Kewen Xiong, Xin Wang

## Abstract

Recent studies have investigated that the synthetic progestins may lead to health issues to the aquatic organisms. Progesterone is a steroidal progestin and has been used as a contraceptive drug, which is detected in the aquatic ecosystem. However, the potential effects of progesterone on the fish reproduction are largely unclear. Here, we tested the effects of progesterone on the fish reproductive and endocrine systems. Adult zebrafish were exposed to progesterone for 10 days at environmental concentrations. The production of eggs was reduced in the exposed fish, and the circulating concentrations of estradiol (E2) and testosterone (T) in female fish or 11-keto testosterone (11-KT) in male fish were significantly diminished. Our results suggested that progesterone may cause adverse health effects on fish by disrupting the endocrine system, and short-term exposure to progesterone could overt affect the fish reproduction.

## Introduction

Many studies reported the endocrine-disrupting compounds reduce the fertility, hatching success, impair the immune system and advance the cancer incidence in wildlife species of aquatic organisms [1-7]. The nature and synthetic steroid hormones are detected in the aquatic environments, which can impair the normal endocrine functions of aquatic organisms [8-13]. Recent studies have demonstrated the natural estrone (E1), estrogens E2, estriol (E3) and estrogen 17-α-ethynylestradiol (EE2) present in the surface waters [14-21]. However, the adverse effects of progesterone on aquatic organisms have not been studied yet.

The zebrafish and human genomes share about 70% conserved syntenic fragments and many homologous genes in zebrafish and human show the similar functions and structures [22-25]. These results provide promising perspective using zebrafish for environmental risk assessment, which can be applied for further researches on humans.

Therefore, we aimed to evaluate the effect of progesterone on the reproduction of zebrafish. We measured the concentrations of hormones both in male and female zebrafish to examine the effect of progesterone on the endocrine functions. Our results showed that progesterone adversely impaired zebrafish reproduction in a dose-dependent manner.

## Materials and methods

### Chemicals

Progesterone was purchased from Sigma-aldrich (P0130-25G). Stock solutions of 0.1, 1, and 10 mg/ml were prepared in ethanol and stored at -20 °C. The stock solutions were diluted with distilled water to the work concentrations of 1, 10, 100 and 1000 ng/L in the exposure system. All the other reagents were of analytical grade.

### Fish husbandry

Adult six-month age zebrafish were maintained in charcoal-filtered water (PH 7.0) at 28 °C with a 14:10 light/dark cycle. The fish were fed a combination of dry flake food (Tetra brand, USA) and live brine shrimp.

### Drug treatment

Ten male and female fish were housed in 50-L glass tanks with 30 L water. Three replicate ranks were used for each group. During the 10-day exposure period, the zebrafish were exposed to nominal concentrations of progesterone (0, 1, 10, 100 and 1000 ng/L), and water was changed daily. The embryos were collected daily and recorded. The zebrafish were raised in accordance with guidelines from the Institutional Animal Care and Use Committee (IACUC) of Huazhong University of Science and Technology.

### Hormone measurement

The blood samples were collected after the progesterone exposure. Plasma extraction and the measurement of sex hormone concentrations were performed as described previously [26]. Briefly, 20 μl plasma were diluted to 500 μl Milli-Q water (Millipore) and extracted with 2 ml ethyl ether twice to collect the ether phase. The concentration of E2, T and 11-KT were detected according to the manufacturers’ instructions (Cayman Chemical Company).

### Statistical analysis

Differences between the control and exposure groups were evaluated by one-way analysis of variance (ANOVA) followed by Tukey’s test. P<0.05 was considered statistically significant.

## Results

### Progesterone exposure decreased the egg productions of zebrafish

Progesterone exposure had no obvious effect on the mortality in all groups during the 10-days treatment. There were no overt differences in the hatching and survival rates between each group, and the malformation rates were similar to the control group (Data not shown).

Embryos production was consistent in each group during the exposure period, the lower concentrations of progesterone (1 and 10 ng/L) appeared no significant impairment to the fish reproduction. When exposed to 100 and 1000 ng/L of progesterone, female zebrafish produced fewer embryos compared to the non-exposed control group (Figure 1).

**Figure 1.**
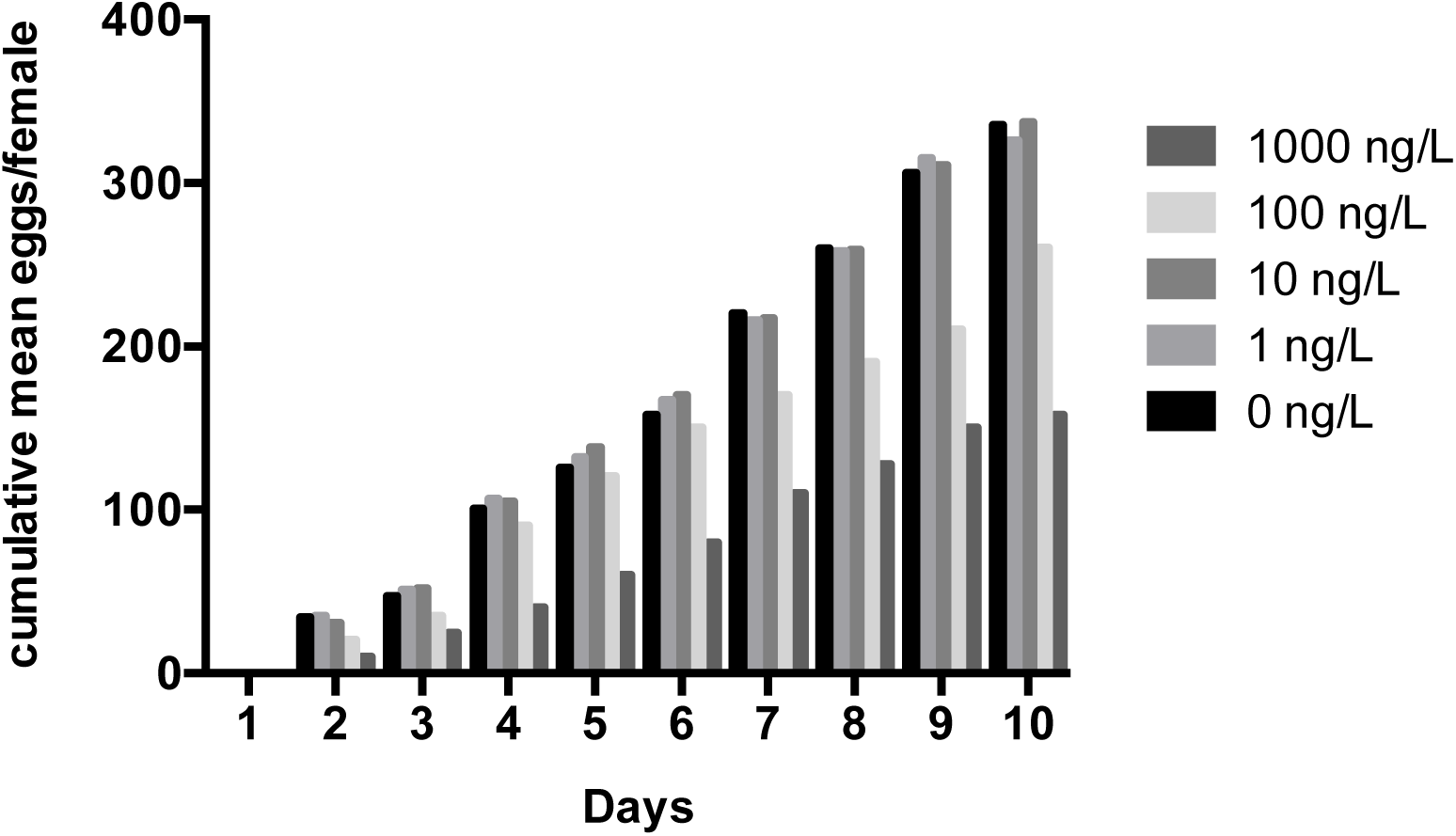
Exposed to progesterone impaired reproduction in zebrafish.

### Progesterone treatment reduced the sex hormone levels in zebrafish

The plasma E2 levels in the female fish were significantly diminished in the 100 and 1000 ng/L of progesterone-treated group (Figure 2). Meanwhile, the T contents were significantly reduced in the 1000 ng/L of progesterone-exposed group (Figure 3). In the male fish, the 11-KT concentrations were significantly decreased when exposed to the 100 and 1000 ng/L of progesterone (Figure 4).

**Figure 2.**
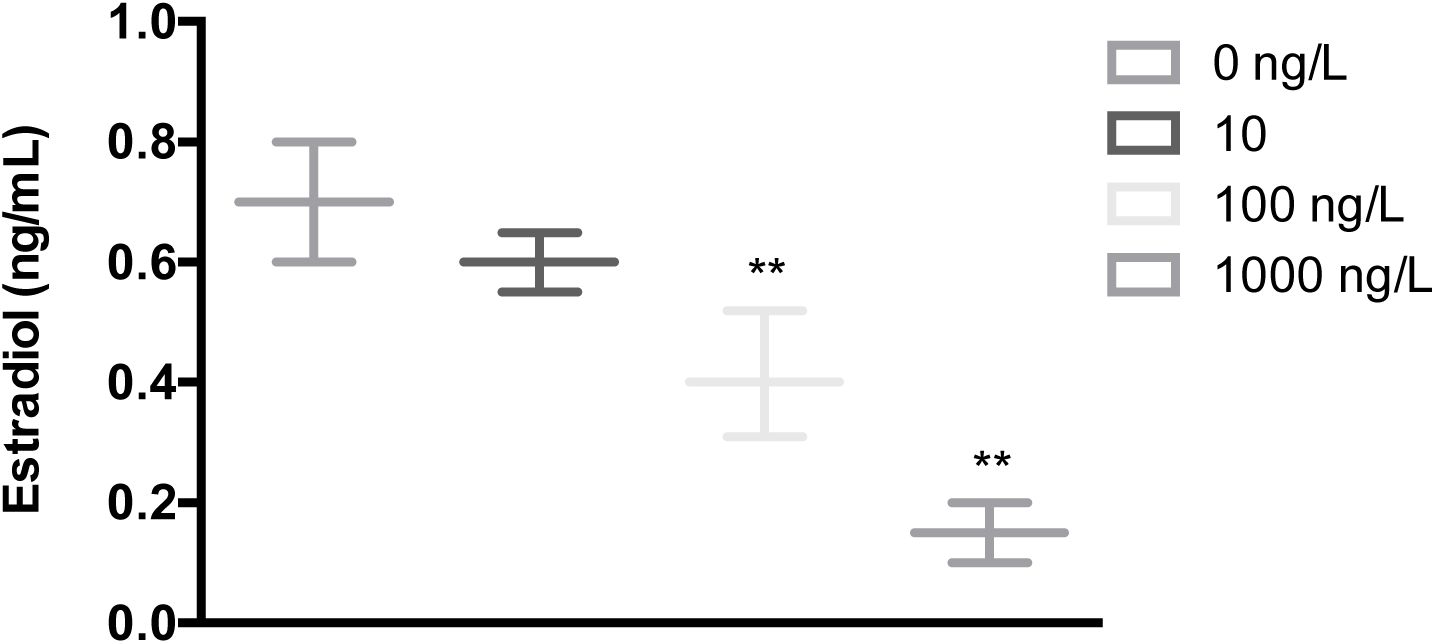
Exposed to progesterone decreased the estradiol (E2) concentrations in female zebrafish.

**Figure 3.**
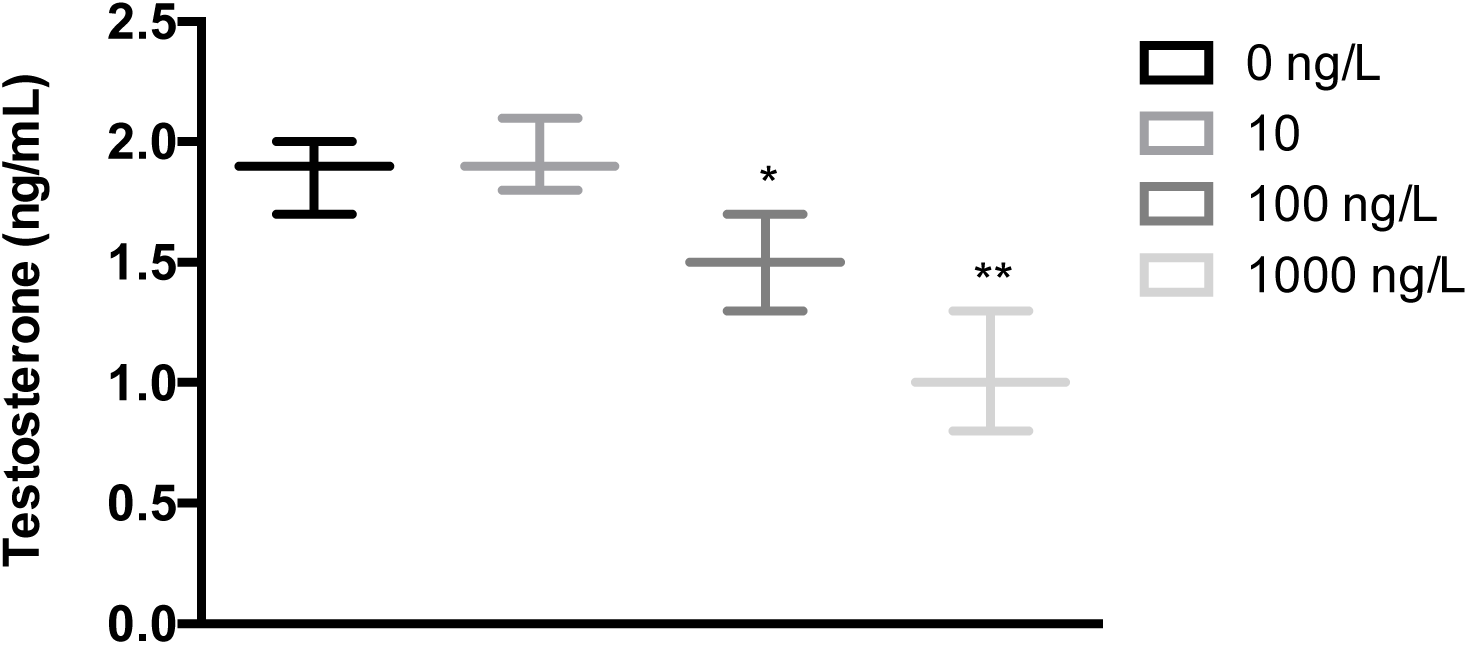
Exposed to progesterone decreased the testosterone (T) concentrations in female zebrafish.

**Figure 4.**
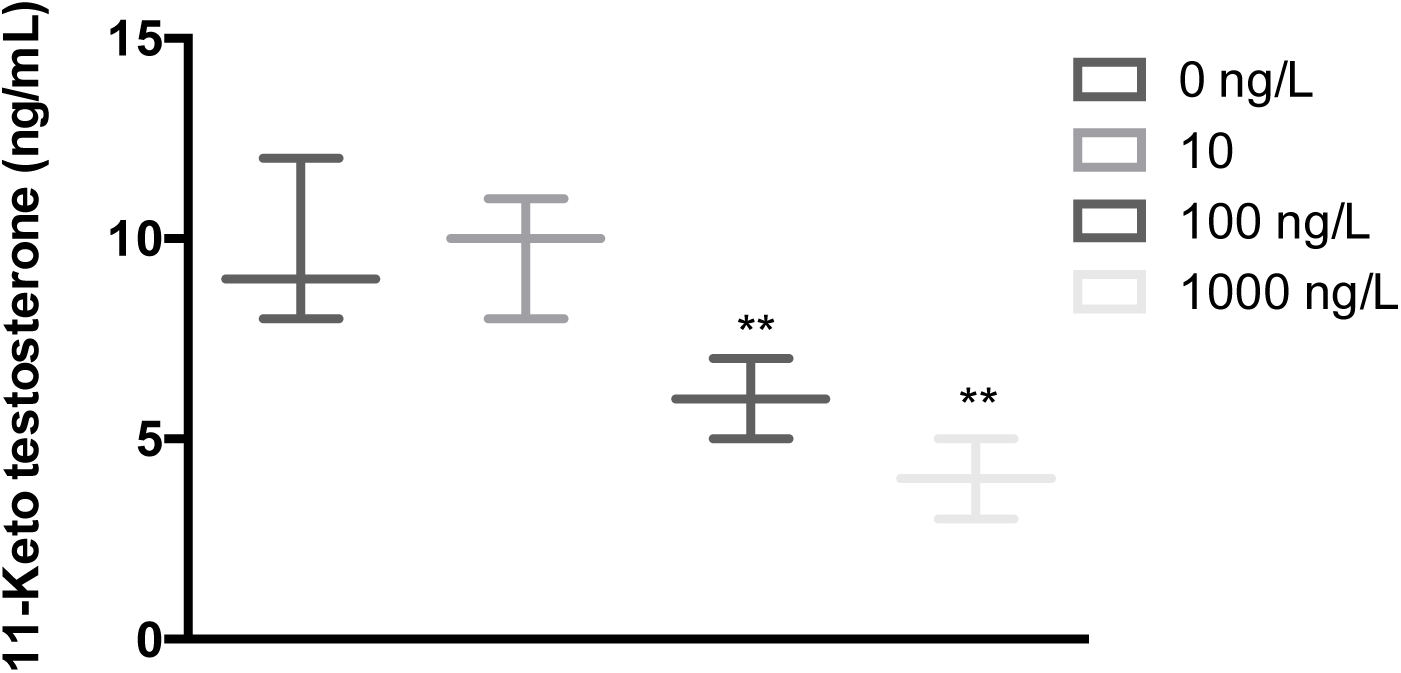
Exposed to progesterone decreased the 11-keto testosterone (11-KT) concentrations in male zebrafish.

## Discussion

Recently, several studies report that natural progesterone may result in endocrine disruption and affect the reproductive system in fish [4, 27-33]. However, the toxicological effects of the progesterone on zebrafish have not been evaluated yet. In this study, our results showed that exposure to progesterone could impair zebrafish reproduction in a short period of time. We also found that the E2 and T levels in female fish and the 11-KT levels in male fish were significantly reduced when treated with progesterone.

Previous studies have demonstrated that progestins have no effect on embryonic development (hatching and survival) [33-36]. Our data are consistent with those previous findings, suggesting that progestins have no obviously acute toxicity to fish eggs. Thus far, the underlying mechanisms of the inhibition of progestins on fish productive system remain unknown. Progestins might mediate the reproductive signaling through progesterone receptors, which might play important roles in the fish.

Our results indicated that exposure to environmental concentrations of progesterone could interrupt the reproductive system in zebrafish. Our findings will be useful to predict the environmental health effects of progesterone on the fish species.

